# Accelerated Discovery of Cell Migration Regulators Using Label-Free Deep Learning-Based Automated Tracking

**DOI:** 10.1101/2025.04.01.646705

**Authors:** Tiffany Chu, Yeongseo Lim, Yufei Sun, Denis Wirtz, Pei-Hsun Wu

**Author notes:** **Corresponding authors:** Pei-Hsun Wu and Denis Wirtz.

## Abstract

Cell migration plays a key role in normal developmental programs and in disease, including immune responses, tissue repair, and metastasis. Unlike other cell functions, such as proliferation which can be studied using high-throughput assays, cell migration requires more sophisticated instruments and analysis, which decreases throughput and has led to more limited mechanistic advances in our understanding of cell migration. Current assays either preclude single-cell level analysis, require tedious manual tracking, or use fluorescently labeled cells, which greatly limit the number of extracellular conditions and molecular manipulations that can be studied in a reasonable amount of time. Using the migration of cancer cells as a testbed, we established a workflow that images large numbers of cells in real time, using a 96-well plate format. We developed and validated a machine-vision and deep-learning analysis method, DeepBIT, to automatically detect and track the migration of individual cells from time-lapsed videos without cell labeling and user bias. We demonstrate that our assay can examine cancer cell motility behavior in many conditions, using different small-molecule inhibitors of known and potential regulators of migration, different extracellular conditions such as different contents in extracellular matrix and growth factors, and different CRISPR-mediated knockouts. About 1500 cells per well were tracked in 840 different conditions, for a total of ~1.3M tracked cells, in 70h (5 min per condition). Manual tracking of these cells by a trained user would take ~5.5 years. This demonstration reveals previously unidentified molecular regulators of cancer cell migration and suggests that collagen content can change the sign of how cytoskeletal molecules can regulate cell migration.

## Introduction

Cell migration is a fundamental and complex biological process that regulates various functions, including immune surveillance and response ^1,2^, embryonic development ^3–5^, and wound healing ^6–9^. Dysregulated cell motility contributes to multiple pathological conditions, such as chronic wounds, fibrosis, and hyper-inflammation^10^. In oncology, motility plays a crucial role in tumor progression, allowing spreading of cancer cells from the primary tumor and their dispersal to distant sites. Identifying the biophysical and molecular regulatory principles that drive cell migration in both healthy and disease contexts have led to the discovery of therapeutic targets^11–13^.

Numerous cell migration assays have been developed to investigate the mechanisms and regulators of cell migration across diverse biological settings. Among these, video-based cell tracking has emerged as a powerful method to study cell migration, as it provides single-cell resolution and temporal data to characterize and understand complex migration processes^14,15^. Tracking cell movements in videos remains a non-trivial task. A critical step in cell tracking is the accurate identification of nuclei/cells. Fluorescent labeling of cells is a common strategy to facilitate cell identification via image processing or machine vision algorithms. However, fluorescence imaging presents several challenges, including phototoxicity and alterations of cell physiology^16–19^. In contrast, brightfield imaging perturbs cellular systems minimally, but detecting cell locations in these images is challenging due to low contrast and highly heterogeneous cell morphology. Moreover, while manual tracking can be used for brightfield videos^16–19^, the extensive effort and subjectivity involved make it impractical.

Traditional cell tracking methods suffer from extremely low throughput, restricting the comprehensive study of cell migration at the molecular level^14,15^. Using conventional manual tracking software, an experienced user may track 50 cells in a video of 50 frames in ~1 hour. Hence, to track cells in 500 different conditions (~1,500 cells per condition) would take a staggering 6 years. Moreover, emerging evidence highlights the context-dependent nature of motility regulation^20,21^, emphasizing the need for large-scale, network-based analyses to fully elucidate the regulatory roles of various molecules and biological context. Evaluating motility responses using existing molecular and drug compound libraries could be highly helpful for identifying novel motility regulators and potential therapeutic targets. However, the lack of high-throughput motility analysis workflows continues to pose a significant challenge. Collectively, these limitations underscore the pressing need for high-throughput motility analysis of unlabeled cells to advance our understanding of cell migration and its intracellular and extracellular mechanisms of regulation.

In this work, we developed a high-throughput, label-free cell motility analysis platform called Deep learning Brightfield Imaging and cell Tracking (DeepBIT). Utilizing convolutional neural networks (CNNs), DeepBIT detects the nuclei of live cells in brightfield images, enabling automated and accurate cell tracking without the need for fluorescence labeling. A common challenge in training CNN models is the need for large, diverse, and representative datasets. We address this by integrating brightfield imaging, molecular labeling, and fluorescence microscopy to generate extensive and varied training datasets with ground truth labels — eliminating the need for time-consuming manual annotations. DeepBIT can track thousands of cells across ~100 time frames in minutes, significantly enhancing the throughput of motility analysis. Its label-free imaging and automated tracking capabilities enable the large-scale deployment of motility studies for drug screening, systems analysis, and high-dimensional interaction studies.

## Results

### Deep learning brightfield imaging and cell tracking (DeepBIT) system

Fluorescently tagged cells or nuclei enable easy and accurate detection of cell and nuclear locations, facilitating cell tracking using real-time imaging. However, fluorescent labeling can potentially alter the biological state of the observed cells, and present issues such as phototoxicity and photobleaching, which limit the duration and temporal resolution of observations. In contrast, brightfield microscopy offers a label-free approach for live-cell imaging with minimal effects on the cells, allowing for extended observation time. However, accurately detecting cell locations in brightfield images is a challenging task^16–18^.

Here, we established a pipeline that accurately detects the locations of label-free cells in brightfield images using convolutional neural networks (CNN), enabling high-throughput cell tracking and analysis (**Fig. 1**). Modern microscopy setups allow for the examination of motility patterns (i.e. trajectories) of individual cells from a vast number of different cell conditions in a 96-well layout, tracking cells over 4 mm^2^ of area in each well with a sufficient temporal resolution of 10 minutes. Combined with high-throughput label-free analysis, our proposed workflow offers the opportunity to study a vast array of motility conditions and perturbations, such as compound screening, factorial extracellular designs, and molecular perturbations, thus enabling systems-level analysis of cell motility.

**Figure 1.**
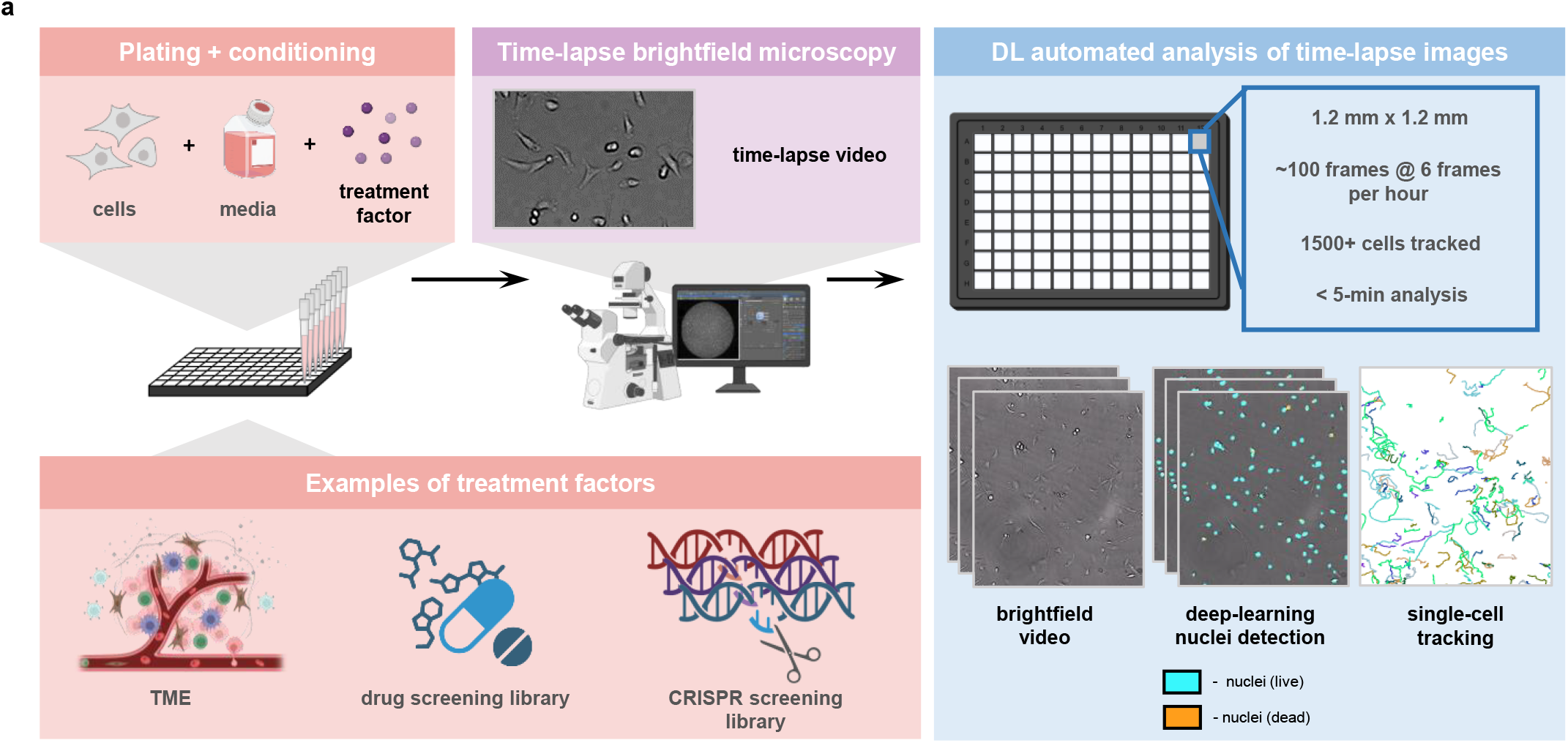
Workflow for plating/cell treatment, DL analysis, and DL model training. **a**. Cells were seeded at 1000-2000 cells/well in 96-well plates and incubated at cell culture conditions (37°C and 5% CO_2_). Following incubation, treatment conditions were added to each well accordingly. The plates were then placed on a microscope (Nikon TI) with an on-stage incubator to maintain cell culture conditions. Time-lapse brightfield images were acquired at 10-minute intervals over a minimum of 16 hours (4 positions per well; 384 positions per plate). The live nuclei in the brightfield images were detected using a custom DL model with DeepLabv3+ network architecture, and the trajectories of individual live nuclei were tracked using the previously established tracking methods.

We established an effective workflow for training a CNN model to detect nuclei locations in brightfield images, bypassing the need for time-consuming manual annotations of training datasets. Molecular labeling with Hoechst 33342 and propidium iodide (PI) was used to label live and dead cells, respectively. Both brightfield and fluorescent images of the cells were acquired simultaneously (**Fig. 2a**; **Supp. Fig. 1**). The live and dead cell map (i.e. ground truth labels) corresponding to the brightfield images was then generated by processing the fluorescent images (**Fig. 2a**). To ensure that robust cell detection across various focal planes was achieved by the trained CNN, a z-stack of 11 brightfield images, representing both in-focus and slightly out-of-focus positions (± 20 µm in z), was acquired at a given field of view (FOV), along with live and dead cell imaging (**Fig. 2b**).

**Figure 2.**
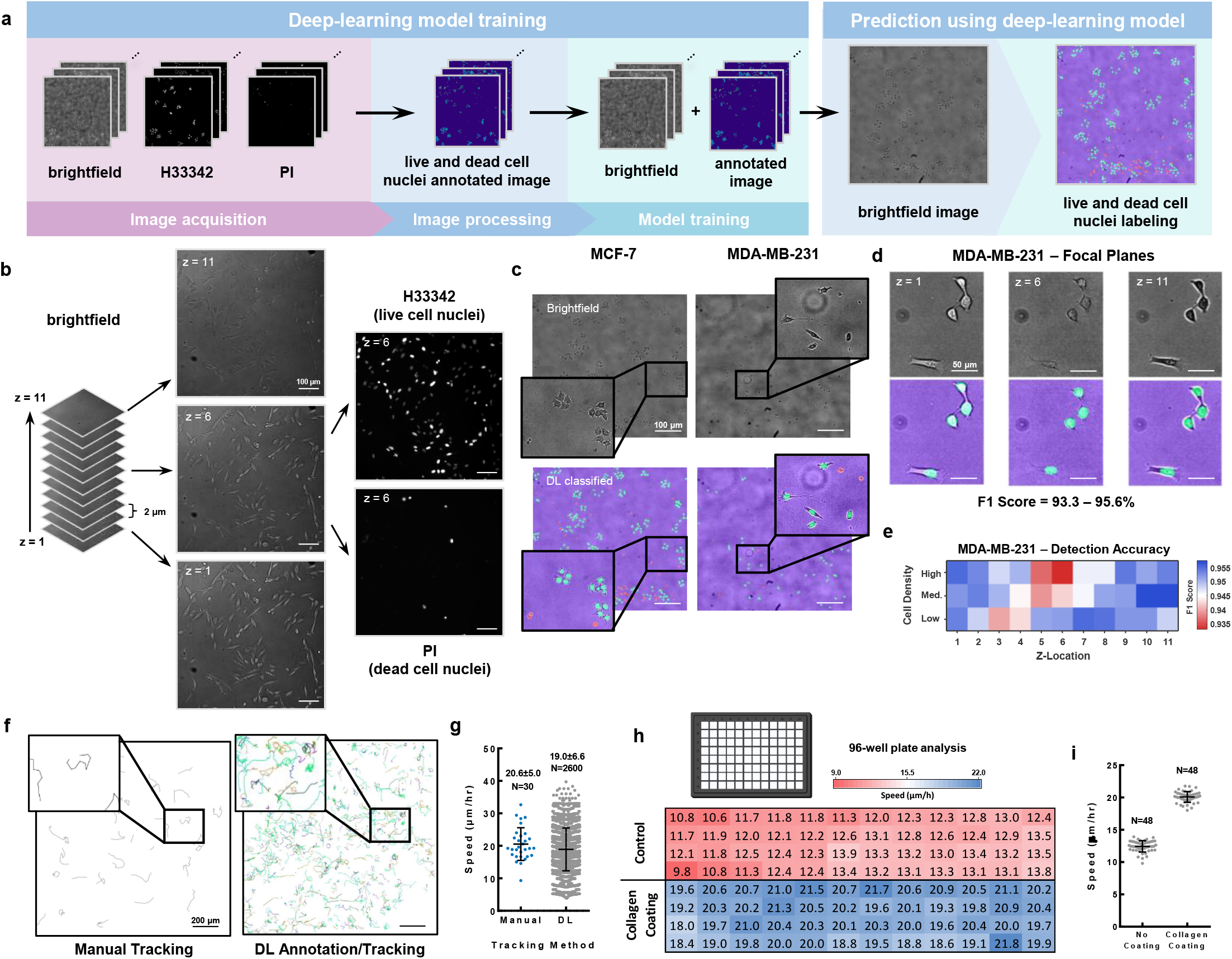
Validation of DL model to ensure high overall accuracy for nuclei prediction. The developed DL model shows a high overall pixel accuracy and can quickly track 1000+ cells in the time it takes to track 30 cells manually. **a**. Training data sets consisting of various cancer cell lines were used to allow for the model to learn to distinguish between live and dead nuclei. Brightfield images with corresponding fluorescent images (H33342 for live cell detection and PI for dead cell detection) were acquired using the same microscopy setup as (**Fig. 1a**). Corresponding brightfield and fluorescent images were combined to generate images annotated for background, live cell nuclei, and dead cell nuclei. These annotated images made up the data used to train the DL model, and the final model was used to annotate brightfield images for live and dead cell nuclei without and use of any additional markers/tags. **c**. Nuclei prediction by the DL model. The brightfield images are at the top, and their respective nuclei predictions by the DL model are on the bottom. Brightfield image of MCF-7 and MDA-MB-231 and the corresponding DL predicted nuclei are present. **d**. Nuclei detection remains robust, and accuracy is retained even at different focal planes. **e**. Nuclei detection remains robust, and accuracy is retained even at different cell densities. **f**. Trajectory comparison between manual tracking (N=30) and DL annotation/tracking (N=2600). Results come from tracking done on the same sample video. **g**. Comparison between individual cell speeds determined from manual tracking (N=30) and DL annotation/tracking (N=2600). Results come from tracking done on the same sample video. **h**. Measured cell speed using DeepBIT for MDA-MB-231 cells in a 96-well plate, with half coated using 50 μg/mL collagen. Measurements remain consistent throughout all replicates within the same plate. **i**. Comparison between MDA-MB-231 cell motility with and without collagen coating. Measurements within the same 96-well plate remain consistent.

This molecular imaging integration allowed us to collect a large dataset of images from both MDA-MB-231 and MCF-7 breast cancer cells — two ubiquitously used cell lines in cancer research — at various cell densities, totaling 1,584 brightfield images across 144 FOVs and 11 focal planes to train (N=792) and test (N=792) CNN models for cell detection (**Fig. 2b**). Live and dead nuclei images were acquired for each FOV to obtain the nuclei labels. The data set is composed of cells at densities of 500 cells/well, 1,000 cells/well, and 2,000 cells/well in a 96-well plate. The molecular images were first converted to the label maps of live and dead cells through image processing. We trained the CNN model with the DeepLabV3+ framework^19^, converting brightfield images into labeled live and dead cell images. The trained model labeled nuclei in the brightfield images of the testing dataset with an overall pixel-level accuracy of 98%+ for both cell types (**Fig. 2c**). The detection of live cells in labeled images showed a robust accuracy with an F1-score of 93.3-95.6% for MDA-MB-231s at different focal planes and cell densities. Interestingly, the cell detection had a better accuracy when images of cells were slightly out of the focal plane (**Fig 2d; Supp. Fig. 2a**). Similarly, the accuracy of the detection of MCF7 cells was robust across focal planes and cell densities, ranging from 90.7%-96.3% (F1-score) (**Supp. Fig. 2c**). Our trained model outperformed cell detection using the pre-trained cyto3 Cellpose model^22^, for which the F1-score was just 76.83% on average for MDA-MB-231 cells and 70.41% for MCF-7 cells (**Supp. Fig. 2d**).

Using our trained CNN, cell locations in each frame of live-cell brightfield movies could be effectively detected to reconstruct cell trajectories (**Fig. 2f**; **Supp. Video 1 & 2**). We further validated the tracked cell speed against manual tracking results (**Fig. 2f, g**). Results showed that the average cell speed measured using DeepBIT (19.0±6.6 μm/h; N=2600 cells) was consistent with the expected values from manual tracking (20.6±5.0 μm/h; N=30 cells). For a single well of a 96-well plate, >2000 cells can be tracked in 100 frames (16 h) within minutes using DeepBIT. The automated tracking throughput is substantially faster compared to manual tracking, which we estimated takes ~1 hour to track 50 cells in 50 consecutive frames. We further demonstrated that cell speed measurements across replicate wells demonstrated high consistency from DeepBIT tracking (**Fig. 2h & i**). Overall, these results demonstrate that the established workflow can accurately analyze the motility of individual cells at high throughput.

### DeepBIT enables the screening of motility regulation compounds

To demonstrate the utility and effectiveness of the DeepBIT workflow, we aimed to identify potential molecular modulators of cell motility by using compounds that inhibit cancer invasiveness and motility using FDA-approved compound libraries. A total of 96 FDA-approved compounds targeting more than 16 distinct signaling pathways (**Supp. Table 1**) were tested on MDA-MB-231 breast cancer cells at three different concentrations (0.01 µM, 1 µM, 10 µM) (**Fig. 3a**). Of the 288 total conditions (96 unique compounds × 3 concentrations), our trained CNN model accurately determined nuclei locations and tracked cell motility under most conditions, successfully analyzing 280 conditions (**Fig. 3b & c**). Eight compounds (**Supp. Table 2**) at a concentration of 100 μM could not be accurately tracked due to the presence of a significant number of particle-like objects in the brightfield images, resulting from the limited solubility of these compounds (**Supp. Fig. 3**). In total, more than 600,000 cells were tracked across the analyzed conditions.

**Figure 3.**
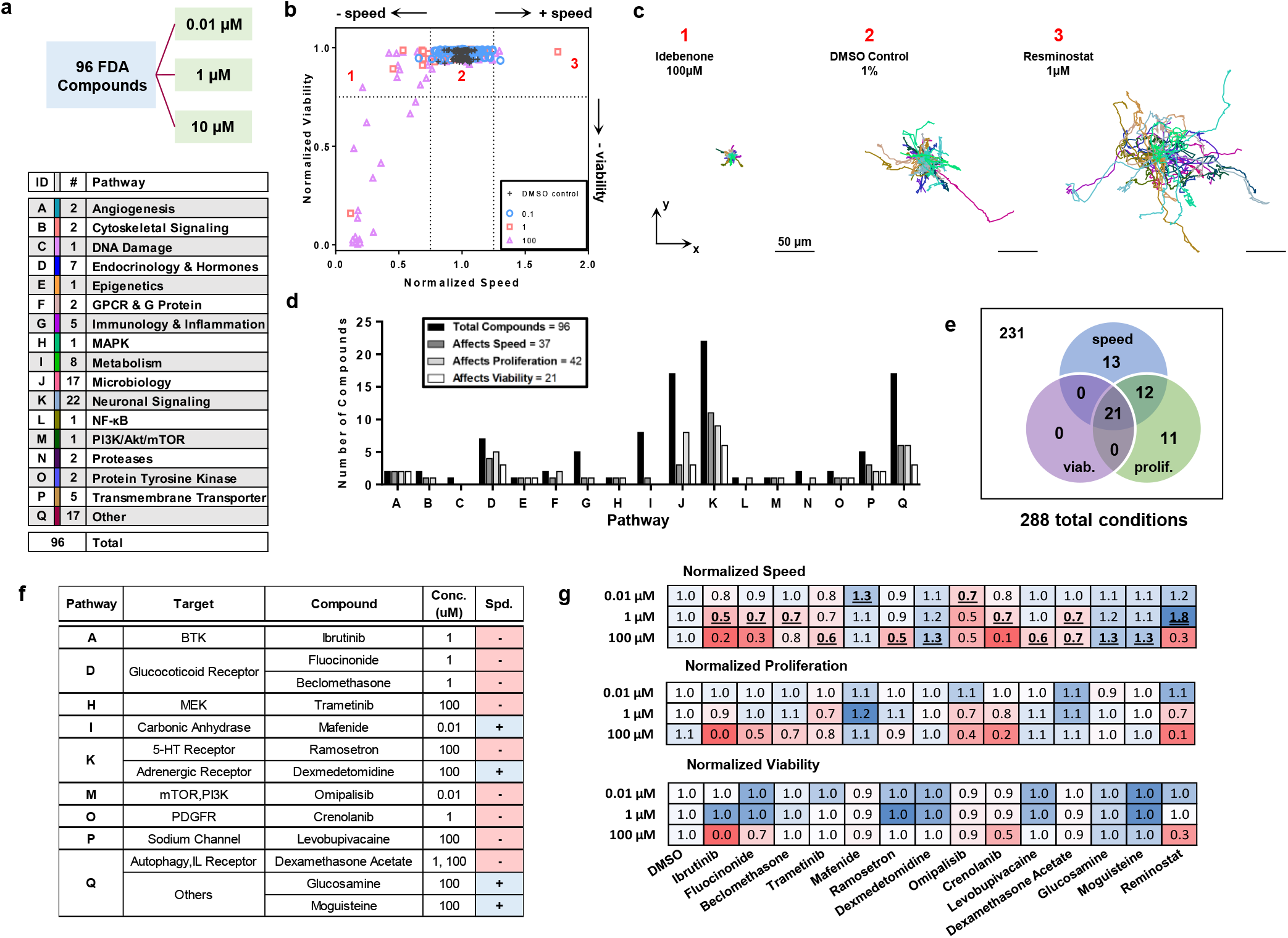
The cell motility response of MDA-MB-231 to a panel of 96 FDA-approved compounds. **a**. Schematic of the compound screening workflow. Cells were incubated with three concentrations (0.01, 1, 100 μM) of compounds for 24 hours prior to imaging. The panel of 96 FDA-approved compounds was categorized into 16 unique pathways. **b**. Comparison between normalized cell speed and viability following treatment for each condition. Dotted lines represent cutoffs for “effective” conditions, and red numbers indicate notable conditions: (1) Idebenone at 100 μM, (2) 1% DMSO; control, and (3) Resminostat at 1 μM. **c**. Representative cell trajectories for visualization of a (1) decrease, (2) control, and (3) increase in cell motility. Numbers correspond with highlighted conditions from panel (b). Scale bar: 50 μm. **d**. Analysis of each pathway, showing the number of compounds which affect motility (black), proliferation (grey), and viability (white) within each pathway category. **e**. Venn diagram that summarizes which conditions significantly affect cell speed, proliferation, and viability out of the total 288 conditions tested (includes all compounds/concentrations). **f**. Table listing the 13 conditions which caused a significant increase (blue, +) or decrease (red, −) in cell speed without influencing proliferation or viability. For each condition, the compound name, pathway, target, and effective concentration is listed. **g**. Heatmaps which show the normalized effect of the 13 compounds from panel (f) on cell speed, proliferation, and viability. DMSO and Reminostat were included as a control condition and significant motility promoter, respectively. Red indicates inhibition and blue indicated enhancement, relative to WT controls.

To ensure that these inhibitors only affected cell motility and not cell viability and proliferation, our live-dead assay was performed for all compounds at each dose. A condition was defined as having an effect on motility, proliferation, or viability if it caused a 25% or greater change compared to the DMSO controls (**Fig. 3b & c**). Among all the tested compounds, we found that 32 affected motility, 42 affected proliferation, and 21 affected viability (**Fig. 3d; Supp. Table 3**). Interestingly, a high occurrence of motility inhibitors was found in the Neuronal Signaling (N=10) and Endocrinology & Hormone (N=4) pathways (**Fig. 3d**). In particular, we found that 5-HT receptor antagonists significantly affected cell motility (**Supp. Fig. 4**).

Since motility inhibition could result from cell killing (and dead cells cannot actively move), we examined the association between cell motility, proliferation, and viability among motility inhibitors. Out of the 46 total conditions that influenced motility, we identified 13 conditions that specifically affected motility without significantly impacting proliferation (**Fig. 3e**). Thus, a majority of conditions that affected motility also affected either proliferation or viability. For the 5-HT receptor antagonists, we found only 1 of the 5 compounds affected only motility (**Fig. 3f**). Not surprisingly, we found that all the compounds that affected viability (N=21) also affected motility and proliferation. We also identified 12 conditions that affected both proliferation and motility.

From the 13 compounds that affected motility only, we found 4 compounds that promoted motility while 9 compounds inhibit motility (**Fig. 3f & g**). Interestingly, we found that Resminostat, an epigenetic drug, could induce an ~80% increase in MDA-MB-231 cell motility at a dosage of 1 μM compared to DMSO controls without affecting viability, but slightly lowered proliferation. However, Resminostat caused complete cell death at 100 μM and had no significant effect at dosages lower than 1 μM (**Fig. 3c & g**). Overall, our results demonstrate that our DeepBIT workflow can efficiently screen libraries of compounds and identify novel potential regulators for cell motility.

### DeepBIT enables combinatorial analysis of extracellular regulators of motility

We next used the DeepBIT platform for a systems analysis of potential extracellular regulators of motility by exploring cell motility responses to combinatorial conditions. A subset of microenvironmental factors known to individually influence breast cancer cell motility were investigated, including cytokine stimulation (EGF, TNF-α)^21^, extracellular matrix (collagen)^23^, and variations in serum (FBS)^24^ (**Fig. 4a**). Additionally, to determine cell-dependent responses, five breast cancer cell lines of varying invasive potential were examined (**Fig. 4a, b**) including MDA-MB-231, MCF-7, SUM149, SUM159, and HCC1954 cancer cells. These cells were chosen because they are routinely used for cancer modeling *in vitro* and in animal models.

**Figure 4.**
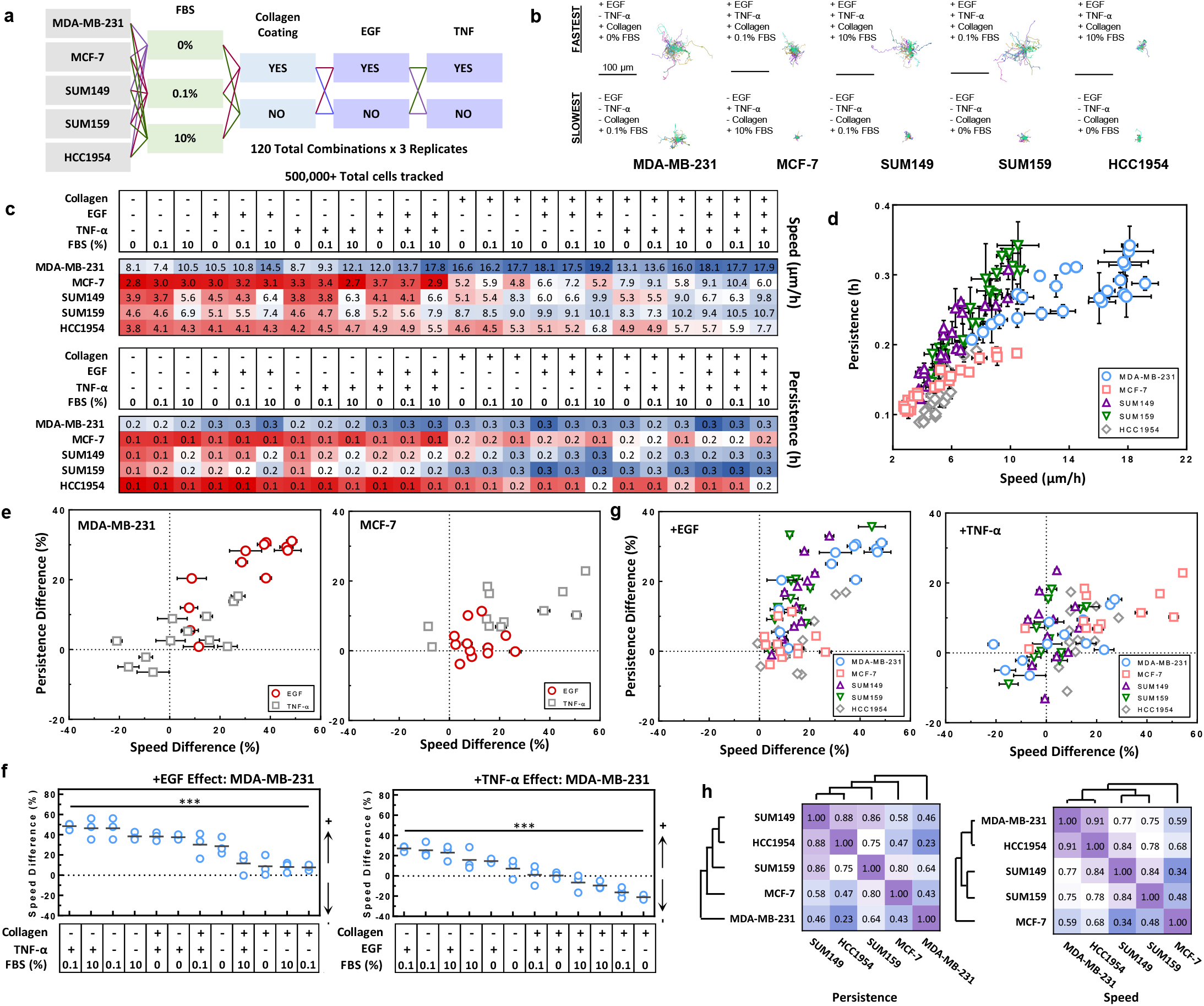
The cell motility response of a breast cancer cell panel to combinations of serum concentration, ECM coating, and cytokines. **a**. Workflow diagram outlining the total unique conditions tested within this study. Cells were serum starved for 16-24 hours and then incubated with their respective cytokines (100ng/mL) for 4 hours prior to imaging. 500,000+ total cells were tracked over 120 combinations with 3 replicates. **b**. Representative cell trajectories for visualization of the fastest and slowest condition for each breast cancer cell line. Scale bar: 100 μm. **c**. Heatmap summarizing the motility (top) and persistence (bottom) effects from the various combinations of microenvironmental factors on breast cancer cells. **d**. Comparison between the persistence and displacement values across all conditions and cell lines. Each point represents a unique combination of microenvironmental factors. **e**. Plots showing the percentage change in speed and persistence induced by EGF (circles) and TNF-α (squares) in MDA-MB-231 and MCF-7 cells. Each point represents a change caused by a soluble factor compared to a baseline combination of factors without the factor of interest. **f**. Comparison of MDA-MB-231 percentage change in speed in response to EGF and TNF-α across all conditions. Shows the difference in magnitude of response depending on the context in which a soluble factor is introduced. Statistical analysis was performed by one-way ANOVA: *P* < 0.05 (*), *P* < 0.01 (**), and *P* < 0.001 (***). **g**. Plots showing the percentage change in speed and persistence induced by EGF and TNF-α across all cell lines. Each point represents a change caused by a soluble factor compared to a baseline combination of factors without the factor of interest. **h**. Hierarchical clustering analysis of speed and persistence changes among the five breast cancer cell lines. Illustrated are the similarities and differences in response to microenvironmental factors between cell lines.

Our trained DeepBIT platform was able to accurately determine nuclei locations and track cell motility for all the above cell lines and conditions, corresponding to a total of 120 unique conditions (**Fig. 4c**). More than 500,000 cells were tracked across the analyzed conditions in three biological replicates. Examination of the heatmaps summarizing speed and persistence values reveals the important effect of extracellular cues in the modulation of cell motility. Collagen and EGF generally increased motility, while the effect of TNF-α varied across cell lines. There was a direct correlation between speed and persistence observed across all cell lines, which suggests that cells that move faster tend to exhibit more directional movement (**Fig. 4d**). Overall, MDA-MB-231 cells had the highest motility, however, their persistence was more susceptible to modulation by microenvironmental factors compared to other tested cancer cells.

To further evaluate how individual cytokines influence motility, we measured changes in speed and persistence in response to EGF and TNF-α (**Fig. 4e & f**). In MDA-MB-231 cells, EGF generally increased both speed and persistence, but the magnitude varied widely (from +7.7% to +48.6% in cell speed) depending on the microenvironment. For instance, EGF enhanced motility by 48.6% in the presence of TNF-α, 0.1% FBS, and collagen on a flat substrate, but this effect dropped to 11.6% when FBS concentration was increased to 10% (**Fig. 4e & f**). Similarly, TNF-α — typically linked to increased cancer cell motility^20,21,25^ — showed both positive and negative effects on MDA-MB-231 motility, depending on extracellular context. TNF-α generally promoted motility, except in environments containing only collagen and/or FBS, where it had a negative effect on migration (**Fig. 4f**). Together, these results highlight that microenvironmental factors critically shape the regulatory impact of motility signals.

We further explored the motility regulatory effects across different breast cancer cell lines. EGF generally enhanced cell motility in all five cell lines under most conditions. However, a significant negative effect was observed in HCC1954 cells, where EGF reduced motility by 18.9% under 10% FBS without collagen or TNF-α (**Fig. 4g & Supp. Fig. 5**). Consistent with earlier findings, TNF-α exhibited a dual role in motility regulation across all cell lines (**Fig. 4g**). Notably, the negative regulation by TNF-α varied by cell line and context. For example, under no collagen, no EGF, and 10% FBS, TNF-α reduced motility in MCF-7 and HCC1954 cells, while under the same conditions, TNF-α strongly promoted motility in SUM149 cells (**Supp. Fig. 5**). Hierarchical clustering of normalized persistence and speed values revealed which cell lines had similar motility responses to extracellular factors (**Fig. 4h and Supp. Fig. 6**). SUM149 and SUM159 clustered closely, which suggests that they share motility characteristics in response to stimuli. MDA-MB-231 and MCF-7 formed distinct clusters of regulated migration, which suggests that they have unique migration strategies in response to microenvironmental factors compared to other breast cancer cells.

Overall, our results demonstrate that breast cancer cell motility is highly context-dependent — including EGF acting primarily as a promigratory factor and TNF-α having pro- and anti-migratory effects. These findings, made possible thanks to our high-throughput assay, provide new insight into how microenvironmental factors regulate cancer cell migration and could help identify potential targets for therapeutic strategies.

### Combining DeepBIT and CRISPR enables deep profiling of motility regulators under diverse microenvironmental conditions

Lastly, we demonstrated that DeepBIT enables deep phenotypic profiling of molecules that regulate cell migration by incorporating molecular perturbation methods, such as CRISPR. We knocked out RHOA, ARPC2, and CTTN in MDA-MB-231 cells; molecules that regulate the assembly and architecture of actin filament network. We measured their motility responses across 16 distinct microenvironmental conditions. These conditions were defined by a combinatorial framework incorporating cytokine stimulation (EGF, TNF-α), extracellular matrix components (collagen), and serum (FBS) (**Fig. 5a**). In total, > 250,000 cells were tracked across the 64 analyzed conditions in three biological replicates.

**Figure 5.**
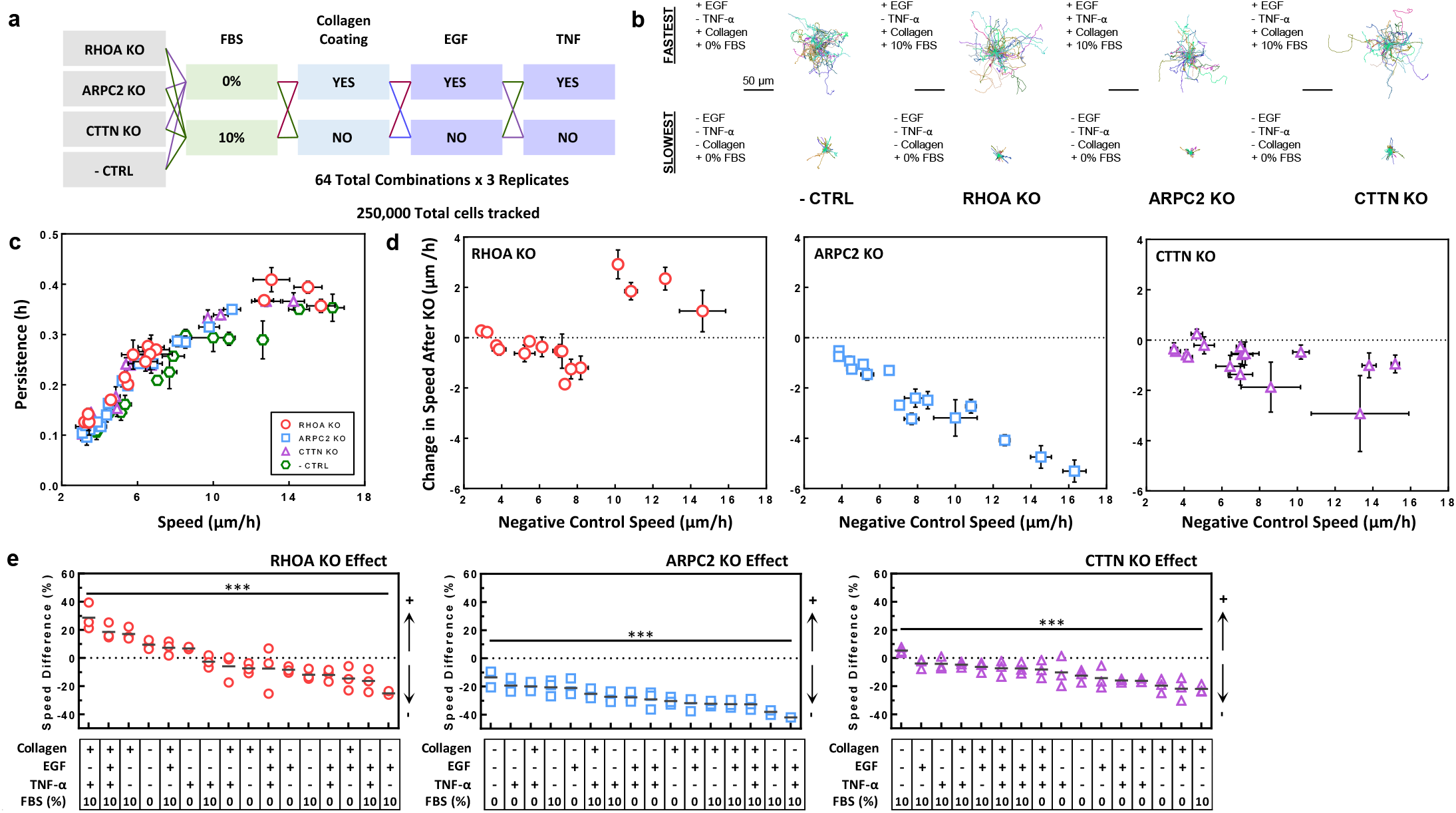
The impact of CRISPR knockouts on cancer cell motility across diverse microenvironmental conditions. **a**. Workflow diagram outlining the total unique MDA-MB-231 knockouts (KO) and conditions tested within this study. KO/WT cells were serum starved for 16-24 hours and then incubated with their respective cytokines (100ng/mL) for 4 hours prior to imaging. **b**. Representative cell trajectories for visualization of the fastest and slowest condition for each unique KO. Scale bar: 50 μm. **c**. Comparison between the persistence and displacement values across all conditions and cell lines. Each point represents a unique combination of microenvironmental factors. **d**. Plots showing the change in speed induced by the KO in MDA-MB-231 cells compared to the negative control speed. Each point represents a unique combination of environmental conditions. **e**. Comparison of percentage change in speed in response to RHOA (circle), ARPC2 (square), and CTTN (triangle) KOs across all microenvironmental conditions. Shows the difference in magnitude of response depending on the context in which KO is introduced. Statistical analysis was performed by one-way ANOVA: *P* < 0.05 (*), *P* < 0.01 (**), and *P* < 0.001 (***).

Molecular knockouts (KOs) can influence cell morphology and we found that RHOA KO induced a notable morphological transformation (**Supp. Fig. 7**). Yet, our trained CNN model continued to successfully detect cell nuclei and tracked motility across all KO experiments, including RHOA KO. Our results revealed that, once more, cell motility under KO conditions exhibited a wide range of behaviors, depending on extrinsic factors (**Supp. Fig. 8**). The slowest motility across all tested KOs and controls consistently occurred in the absence of external stimuli (i.e.: no FBS, collagen, EGF, or TNF-α). In contrast, the highest motility responses for both KOs and controls were observed under conditions with different combinations of three or more external stimuli (**Fig. 5b-e**).

Interestingly, we found a strong association between cell speed and movement persistence across all KO groups and conditions (**Fig. 5c**). At lower speeds, persistence was highly correlated with speed, whereas at higher speeds, persistence plateaued. These results suggest a “universal” motility response characterized by the non-linear relationship between cell speed and persistence^26^.

We further examined the effect of these molecular manipulations on cell motility across different conditions, comparing them to the scramble control. The loss of RHOA induced distinct effects on cell motility that depended on extrinsic conditions. Under conditions where cells migrated at high speed, RHOA KO further enhanced cell migration. In contrast, when cells were already in conditions associated with lower migration potential, RHOA KO led to a decrease in motility (**Fig. 5d**). Knockout of ARPC2, on the other hand, generally led to a reduction in cell speed. The extent of this reduction was strongly associated with the cell’s migration capability, with an estimated ~35% decrease in speed observed across all tested conditions (**Fig. 5d**). The mixed effects of CTTN KO in MDA-MB-231 cells were also observed, with most conditions showing minimal decreases in speed. We further examined how extrinsic conditions are associated with different molecular KOs (**Fig 5d**). In the presence of collagen and serum, RHOA KO significantly increased cell speed (**Fig. 5e**). Under other conditions, RHOA KO caused a slight decrease in cell motility or had minimal effects. The ARPC2 KO consistently reduced cell motility across most conditions, with the largest effect observed in the presence of collagen (**Fig. 5e**). Minor reductions in speed were observed for CTTN KO under certain conditions (**Fig. 5e**).

Our findings demonstrate that motility regulation via gene targeting is also context-dependent. ARPC2 is clearly essential for maintaining migration speed, whereas RHOA and CTTN appear to have a more context-dependent role. These results, rendered possible by our high-throughput assay, provide insight into how cytoskeletal regulators influence cancer cell motility and suggest targets that could be used to modulate migration.

## Discussion

We demonstrate that the proposed DeepBIT framework enables accurate and efficient tracking of cell motility in label-free brightfield videos using a trained CNN. Our workflow addresses the challenges of training accurate CNN models for nuclei detection by integrating fluorescent cell labeling with brightfield microscopy. This approach generates an extensive training dataset without the need for potentially time consuming and subjective manual annotations. The automated labeling of nuclei in brightfield imaging provides a foundation for high-throughput analysis of cell migration behaviors at a single-cell resolution in a time-resolved manner. Additionally, it facilitates large-scale screening of motility inhibitors and enables the exploration of motility responses at complex molecular intersections. The convolutional neural network (CNN) model developed in this study for live nuclei detection is effective in tracking cells with epithelial or mesenchymal phenotypes, such as MCF-7 or MDA-MB-231 cell lines. However, its performance may be limited when applied to cells with significantly different morphologies, such as lymphocytes. A major challenge in retraining CNN models is the need for substantial datasets for training and testing to ensure robust performance. Our proposed workflow streamlines the acquisition of large training datasets, enabling efficient retraining of the cell detection CNN model to accommodate various cell lines.

The role of RhoA in cancer invasion remains controversial, as findings from individual studies have reported both pro-invasive and inhibitory effects when RhoA is inhibited^27,28^. Our study also demonstrates that RhoA knockout induces a wide range of motility responses, from inhibition to minimal or even enhanced motility, depending on the presence of collagen in the environment. These results underscore the complex interplay between extrinsic and intrinsic cellular factors in regulating motility. A single-axis (piecemeal) analysis of molecular function provides a limited context and may not accurately reflect its role in the complex *in vivo* environment, emphasizing the need for high-throughput approaches to capture a more comprehensive picture and precision of biology.

Automated cell tracking in brightfield images significantly reduces manual effort and enables high-throughput analysis of cell motility, facilitating advances in systems and precision biology. In this study, we analyzed a total of 840 experiments (including all repeats and conditions) with 100+ frames per experiment, requiring approximately 70 hours of processing time and tracking ~1.2 million cells. By comparison, we estimate that manual tracking by a trained researcher achieves a throughput of 50 cells across 50 frames per hour, meaning the same task would take approximately 48,000 hours (~5.5 years) to complete manually. This stark contrast highlights the potential of automated tracking for large-scale motility studies.

Our assay reveals that 5-hydroxytryptamine (5-HT) antagonists of can significantly impact cell motility, implicating neurotransmitter signaling pathways in the regulation of breast cancer migration. While the role of 5-HT receptors, or serotonin receptors, is well established in neurotransmitter regulation, recent studies have also highlighted their involvement in tumorigenesis and tumor progression across various tumor types^29,30^. In breast cancer, overexpression of 5-HT has been observed in patients with triple-negative breast cancer^31^ and is associated with increased pro-tumor activity. Additionally, treatment with 5-HT agonist antidepressants in cancer patients has been linked to a higher risk of cancer recurrence^32–34^. Our findings further demonstrate that a subset of 5-HT antagonists induces anti-migratory effects in cancer cells, independent of their anti-proliferative activities. These results provide further evidence of the role of the 5-HT-associated pathway in regulating cancer invasion.

Our assay identified dose-dependent, conflicting effects of a pan-histone deacetylase (HDAC) inhibitor, resminostat, in triple-negative breast cancer cells (MDA-MB-231). Various HDAC inhibitors have been studied for their anti-tumor effects in breast cancer^35,36^. In particular, suberoylanilide hydroxamic acid (SAHA), a well-studied pan-HDAC inhibitor, has been shown to suppress cancer cell proliferation, migration, invasion, and epithelial-mesenchymal transition (EMT)^37–39^. However, conflicting reports suggest that SAHA may also promote cancer invasion and EMT in breast cancer cells^40,41^. Resminostat, another pan-HDAC inhibitor, has been extensively investigated for its anti-tumor effects in hepatocellular carcinoma and T-cell cutaneous lymphoma^42,43^. However, its effects on breast cancer remain less explored. While high doses (100 µM) of resminostat inhibit breast cancer cell proliferation, exposure to low doses (1 µM) significantly increases cell motility, indicating an enhanced invasive state. Together, our findings highlight the complex, dose-dependent effects of HDAC inhibitors on breast cancer cells, emphasizing the need for further investigation to optimize their therapeutic potential.

## Abbreviations

(DeepBIT): Deep learning brightfield imaging and cell tracking
(FOV): Field of view
(CNN): Convolutional neural networks
(KO): knockout
(PI): Propidium iodide
(EGF): Epidermal growth factor
(TNF-α): Tumor necrosis factor-alpha
(RHOA): Ras homolog family member A
(ARPC2): Actin-related protein 2/3 complex subunit 2
(CTTN): Cortactin

## Author contributions

T.C., Y.L., P.H.W. and D.W. designed the experiments. T.C., Y.L., and Y.S. collected the data; P.H.W., and Y.L. developed the analytical tools and analyzed the data. T.C composed the figures and wrote the supplemental material and supplemental figures. T.C., P.H.W., D.W., and Y.L. wrote the manuscript. T.C, P.H.W., and D.W. edited the manuscript.

## Acknowledgements

The authors acknowledge the following sources of support: UG3CA275681 (PHW), UH3CA275681(PHW); U54AR081774 (DW); U54CA268083 (DW); R01CA300052 (DW), all from the National Institutes of Health.

## Materials and Methods

### Cell lines and culture

MDA-MB-231 (HTB-26), MCF-7 (HTB-22), and HCC1954 (CRL-2338) cells were purchased from the American Type Culture Collection (ATCC), and SUM149 (HUMANSUM-0003004) and SUM159 (HUMANSUM-0003006) were obtained from BioIVT. The breast cancer cell lines were cultured in DMEM supplemented with 10% fetal bovine serum (FBS, Corning, 35-010-CV) and 1% penicillin-streptomycin (P/S, Sigma-Aldrich, P0781-100ML). Cells were maintained at 37°C and 5% CO_2_ in an incubator for passage numbers less than 15. Cell were trypsinized (trypsin EDTA, Sigma-Aldrich, T4049-500ML) and passaged at 70-80% confluency, every 2-3 days.

### Live and dead cell assay and analysis

Cells were plated at a density of 1500-2000 cells per well depending on the cell line. Following an overnight incubation and/or cell tracking experiment, well plates were stained using Hoechst 33342 (Thermo Fisher Scientific, H21492) and propidium iodide (PI, Thermo Fisher Scientific, P1304MP), and imaged in two channels, 395 and 555 nm. Cells were identified in the fluorescent images and were classified as live or dead using filters based on Hoechst 33342 and PI intensity.

### Deep-learning model training

Cells were plated at varying densities of 500, 1,000, and 2,000 cells per well in 96-well plates (Corning, 3603). After plating, cells were incubated overnight to allow attachment to the surface. The following day, cells were stained with Hoechst 33342 (3 µg/mL) and propidium iodide (PI, 3 µg/mL) to label live and dead nuclei, respectively, for 30 minutes. For imaging, brightfield images were first captured at 11 different focal planes, offset above and below the in-focus plane, using a Nikon TI-E microscope with 10x objective. Immediately after brightfield imaging, fluorescence images of Hoechst 33342 and PI were acquired for each field of view (FOV) to minimize potential differences caused by cell motility. Four FOVs were imaged per well.

Live and dead cell-labeled images for each FOV were generated from the fluorescent images. A bandpass filter was applied to reduce background noise and improve the signal-to-noise ratio^44,45^. Intensity thresholds were manually determined to identify positively stained regions. Since Hoechst stains both live and dead nuclei, regions positive for both Hoechst and PI were classified as PI-positive (dead) cells. The brightfield images were also normalized prior to training. First, the background field was estimated and subtracted from each brightfield image. The background field was calculated using a 2D median filter with a window size of 300 × 300 pixels. After background subtraction, the images were rescaled to have a mean intensity of 100 arbitrary units (a.u.) and a standard deviation of 30 a.u. This was achieved by dividing the pixel intensities of the background-corrected images by their standard deviation, multiplying by 30, and then adding 100.

The image set is then split into a training and testing data set to train a DeeplabV3+ network^19^ to detect live and dead cell label from brightfield images. The image split is performed at FOV levels such that the whole z-stack images are assigned either to training or testing. The images are first down-sampled at two-fold and the image size for training models are set to 1024x 1024 pixels. The networks are trained with 30 epochs. The model performance is evaluated based on the testing image. All computational procedures including image processing and model training were performed using MATLAB (The MathWorks, Natick, MA).

### Cell motility assay

Black 96-well plates (Corning, 3603) were coated with 50 μg/mL of collagen type I, rat tail (Corning, 354249) for 30 minutes. Breast cancer cells were seeded at so that cells would reach ~2000 cells per well prior to imaging. To ensure even cell distribution, the 96-well plate was placed on a shaker at 500 RPM for 2 minutes following seeding and allowed to settle for 20 minutes before being placed in an incubator. Cells underwent desired treatments following an overnight incubation. Following treatment, 96-well plates were placed onto a microscope (Nikon Eclipse Ti-E) with an on-stage incubator (Tokai Hit, INU-TIZW) to maintain cell culture conditions of 37°C and 5% CO_2_. Time-lapse brightfield images were acquired at 10-minute intervals over a minimum of 16 hours (4 positions per well; 384 positions per plate). Images were obtained at 10x magnification.

### FDA-approved compound treatment

MDA-MB-231 cells were seeded at 1,000 cells per well in a 96-well plate. Cells were treated with 0.01 μM, 1 μM, and 100 μM FDA-approved compounds randomly selected from a single plate of an FDA-approved screening library (Selleck Chem, L4300-03) for 24 hours. Treated cells underwent time-lapse imaging with subsequent live/dead cell analysis. “Control” samples were supplemented with DMSO vehicle in place of FDA-approved compounds.

### CRISPR lipofection

MDA-MB-231 cells were seeded in 12-well plates so that they would reach 60-80% confluency on the day of transfection. Cells were transfected with TrueCut Cas9 Protein (2500 ng per well, Thermo Fisher Scientific, A36498) and RHOA (CRISPR1031551_SGM), ARPC2 (CRISPR995699_SGM), and CTTN (CRISPR925694_SGM) True Guide sgRNA (480 ng per well, Thermo Fisher Scientific, A35533) using Lipofectamine CRISPRMAX (Thermo Fisher Scientific, CMAX00003) for 48 hours. Following transfection, cells were trypsinized and seeded into 96-well plates for high-throughput cell motility assay conditioning and analysis. Non-targeting sgRNA-treated (Thermo Fisher Scientific, A35526) cells and cells treated with Lipofectamine CRISPRMAX only were used as controls.

### Serum starvation and cytokine treatment

MDA-MB-231 and HCC1954 were seeded at 1,500 cells per well and MCF-7, SUM159, and SUM149 were seeded at 1,250 cells per well in a 96-well plate. Cell culture medium was replaced with starvation medium (DMEM supplemented with 0%, 0.1%, 10% FBS respectively and 1% P/S) following an overnight incubation after plating. Cells were serum-starved for 16-24 hours. 100 ng/mL of EGF (PeproTech, AF-100-15), TNF-α (PeproTech, 300-01A), and EGF+TNF-α was added to each well respectively and incubated with cells for 4 hours prior to imaging. “Control” samples underwent serum-starvation and were supplemented with additional starvation medium in place of cytokines.

### Deep-learning cell detection, tracking, and analysis

To track the cell motility, the live cell nuclei locations in the brightfield images are first detected using the trained DL model after 2-fold down sampling. The marker-controlled watershed was then implemented to segmentation, the detected live nuclei image and locations of the segmented nuclei object are then measured^44^. Objects with area less than 15 um^2^ are excluded from further analysis. Once the nuclei locations are obtained from all timeframes, cell trajectories are tracked using previously established methods^46,47^. Cell instantaneous speed and persistence are calculated for each tracked object. Cell instantaneous speed was calculated at a time-lag (*τ*) of 1 hour using the following equation:

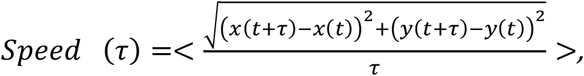

where *τ* represents time-lag, <…> indicates time-averaging, *t* represents the instantaneous time, and (*x,y*) are the coordinates for cell location at a given point in time.

Persistence is defined as the ratio of net distance traveled (D_net_) to integrated distances traveled (D_int_), calculated using the following equations:

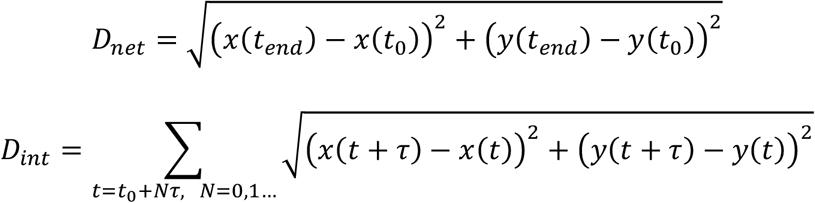

In these equations, *t*_*0*_ represents the initial timepoint, *t*_*end*_ represents the final timepoint collected, and *N* represents the total number of time steps, taking into account the time lag. The time lag for calculating *D*_int_ is a single frame.

### Statistical analysis

One-way analysis of variance (ANOVA) was performed using GraphPad Prism and was used to determine statistical significance. Results were considered significant at *P* < 0.05 (*), *P* < 0.01 (**), and *P* < 0.001 (***).

