## supplementary figures for "Accelerated Discovery of Cell Migration Regulators Using Label-Free Deep Learning-Based Automated Tracking"

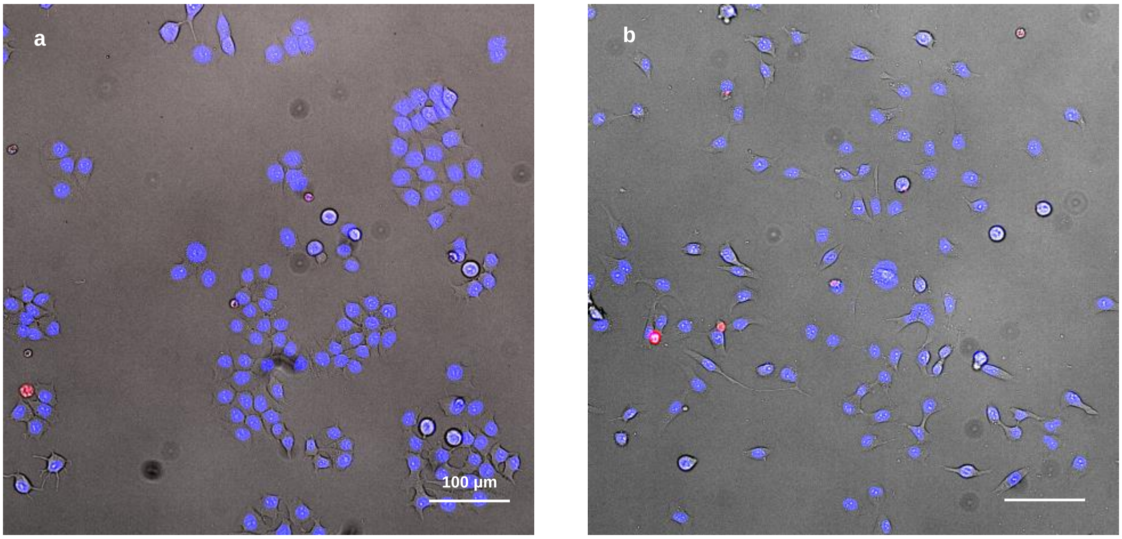


**Supplementary Figure 1. Examples of images used to train the CNN for cell detection in brightfield without the need for molecular labeling or manual annotation.** Representative brightfield image of **(a)** MCF-7 and **(b)** MDA-MB-231 cells with H33342 (blue; live cell nuclei) and PI (red; dead cell nuclei) signal overlayed. The live and dead cell map was used as ground truth labels for the corresponding brightfield image. The brightfield images with corresponding ground truth labels, allowed us to train a highly accurate CNN for cell detection without the use of any additional manual annotation.

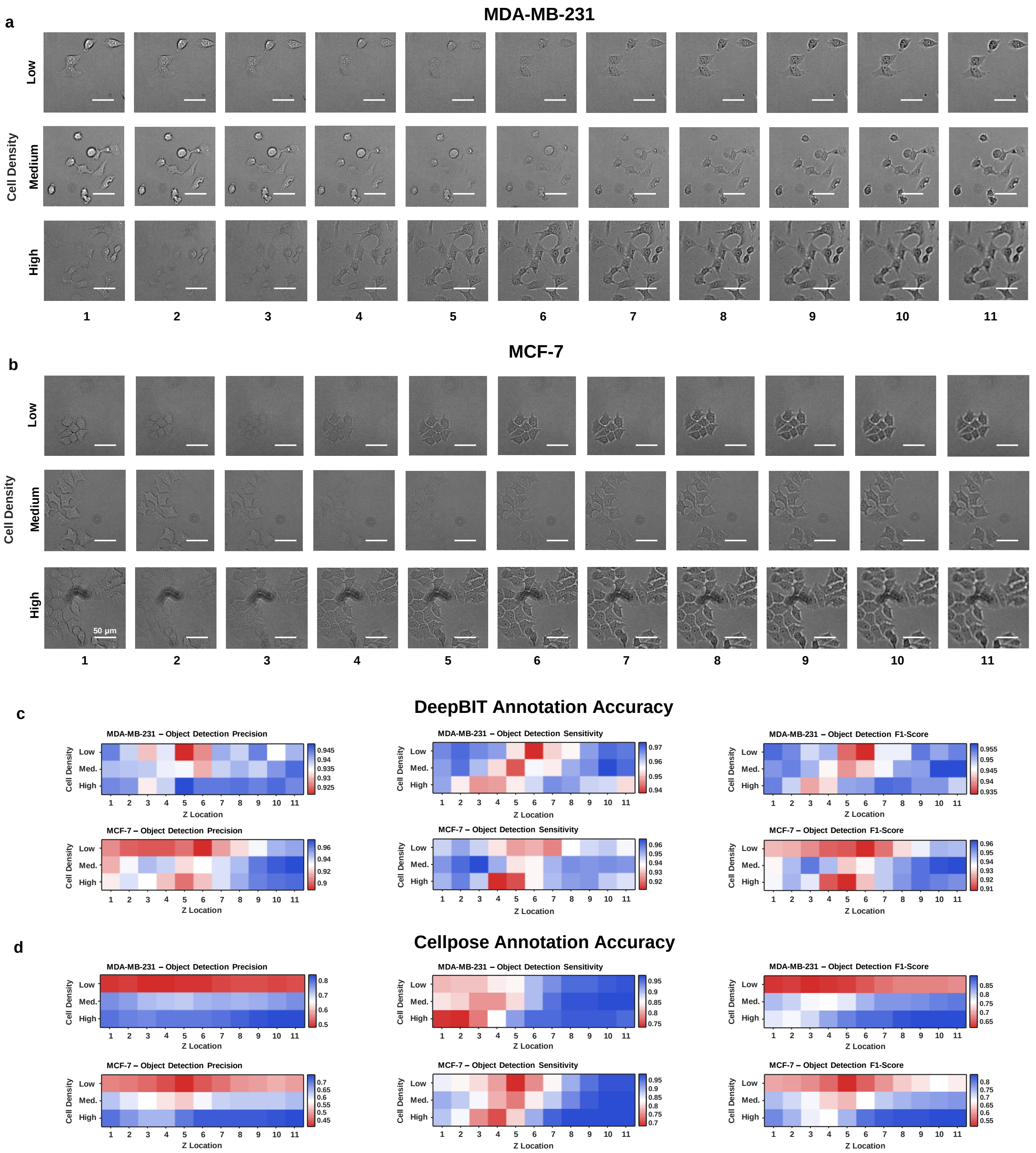


**Supplementary Figure 2. Comparison between cell annotation accuracy of DeepBIT and Cellpose.** Representative brightfield images of **(a)** MDA-MB-231 and **(b)** MCF-7 from the testing data set. This data set is composed of various cell densities including 500 cells/well (low), 1,000 cells/well (med.), and 2,000 cells/well (high). Each FOV was imaged across 11 focal planes with a step size (z) of 2 μM. Accuracy values were obtained using generated cell masks from **(d)** DeepBIT and **(e)** Cellpose (cyto3) compared to the ground truth labels. Accuracy values for object detection precision, sensitivity, and F1-score were calculated for MDA-MB-231 and MCF-7 for all densities and focal planes.

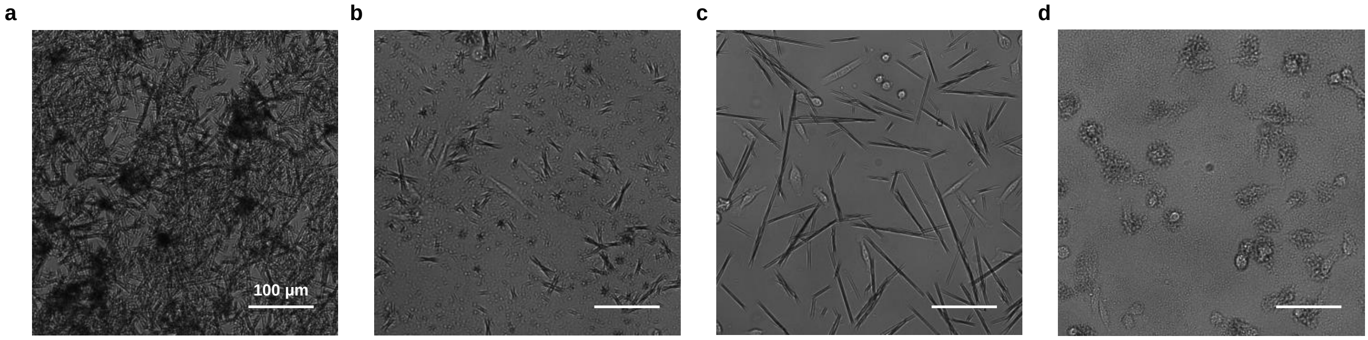


**Supplementary Figure 3. Examples of particle-like objects present in brightfield images which impact cell detection and tracking.** Representative images of 100 µM **(a)** Benactyzine, **(b)** Fluocinonide, **(c)** Atovaquone, and **(d)** Dolutegravir showing large, opaque particles from compounds blocking visualization and analysis of cells. The abundance of particles caused DeepBIT to not be able to detect cells accurately, and, in some cases, it is even difficult to manually detect the cells present in the images. At lower concentrations it was possible to obtain motility results for these compounds using DeepBIT. Some of these particles did not prevent analysis using fluorescence for cell proliferation and viability results.

**a**

| PATHWAY | TARGET | COMPOUND | SPD. | VIAB. | PROLIF. |
|---|---|---|---|---|---|
| Neuronal Signaling | 5-HT Receptor | Sertraline | - | - | - |
| | | Ramosetron | - | | |
| | | Sarpogrelate | - | - | - |
| | | Escitalopram | - | | - |
| | | Flibanserin | - | | +/- |
| | Adrenergic Receptor | Indacaterol | - | - | - |
| | | Vilanterol | - | - | - |
| | | Dexmedetomidine | + | | |
| | | Guanabenz Acetate | | | - |
| | | Buflomedil | | | |
| | AChR | Benactyzine | | | |
| | | Arecoline | | | |
| | | Decamethonium Bromide | | | |
| | | Methacholine | | | |
| | | Oxybutynin | - | N/A | N/A |
| | Histamine Receptor | Doxepin | - | - | - |
| | COX | Nabumetone | | | |
| | | Acemetacin | | | |
| | | Mefenamic Acid | | | |
| | | Diclofenac Diethylamine | | | |
| | P2 Receptor | Ticagrelor | - | - | - |
| | GluR | Piracetam | | | |

**b**

5-HT Receptor Compounds

Norm. Speed

| | DMSO | Sertraline | Sarpogrelate | Ramosetron | Escitalopram | Flibanserin |
|---|---|---|---|---|---|---|
| 0.01 µM | 1.00 | 0.96 | 0.83 | 0.91 | 0.89 | 1.05 |
| 1 µM | 1.00 | 0.98 | 0.88 | 0.92 | 0.87 | 1.04 |
| 100 µM | 1.00 | 0.14 | 0.16 | 0.53 | 0.42 | 0.72 |

Norm. Proliferation

| | DMSO | Sertraline | Sarpogrelate | Ramosetron | Escitalopram | Flibanserin |
|---|---|---|---|---|---|---|
| 0.01 µM | 1.01 | 0.94 | 1.07 | 1.00 | 0.99 | 1.13 |
| 1 µM | 1.01 | 1.02 | 1.05 | 1.07 | 0.99 | 1.37 |
| 100 µM | 1.05 | 0.00 | 0.00 | 0.91 | 0.70 | 0.55 |

Norm. Viability

| | DMSO | Sertraline | Sarpogrelate | Ramosetron | Escitalopram | Flibanserin |
|---|---|---|---|---|---|---|
| 0.01 µM | 0.98 | 0.99 | 0.99 | 0.99 | 0.98 | 0.98 |
| 1 µM | 0.98 | 0.98 | 0.99 | 0.99 | 0.98 | 0.99 |
| 100 µM | 0.98 | 0.01 | 0.01 | 0.98 | 0.97 | 0.85 |

**Supplementary Figure 4. Several compounds associated with neuronal signaling affected MDA-MB-231 motility. (a)** List of tested compounds in neuronal signaling and their associated effect in motility and proliferation. Negative effects are noted in red (-), positive in blue (+), and dosage dependent effect are noted in purple (+/-). All 5-HT receptor antagonists affected motility. **(b)** The Heatmaps show the normalized effects of all 5-HT receptor antagonists on motility, proliferation, and viability at concentrations of 100 µM, 1 µM, and 0.01 µM.

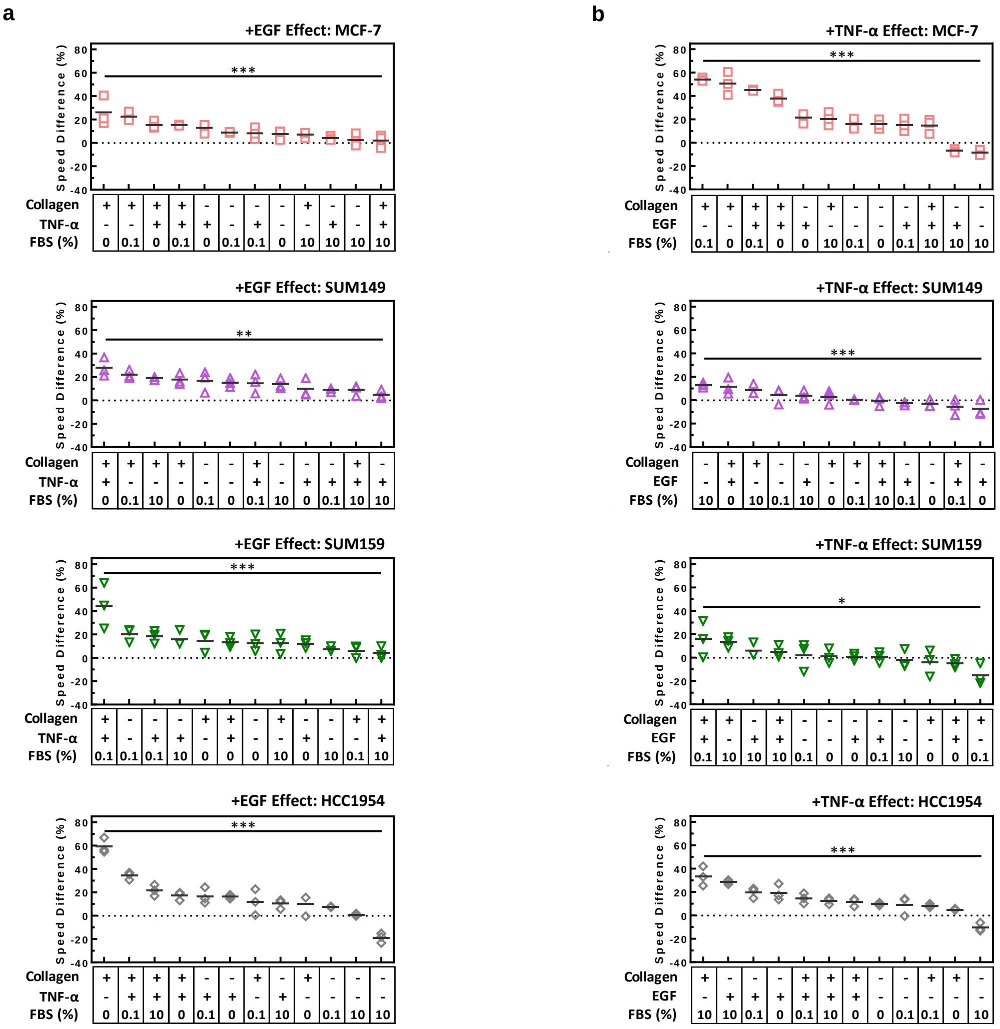


**Supplementary Figure 5. The cell motility response of a breast cancer cell panel to combinations of serum concentration, ECM coating, and cytokines.** Comparison of MCF-7, SUM149, SUM159, and HCC1954 percentage change in speed in response to **(a)** EGF and **(b)** TNF-α across all conditions. Shows the difference in magnitude of response depending on the context in which a soluble factor is introduced. Statistical analysis was performed by one-way ANOVA: $P < 0.05$ (*), $P < 0.01$ (**), and $P < 0.001$ (***).

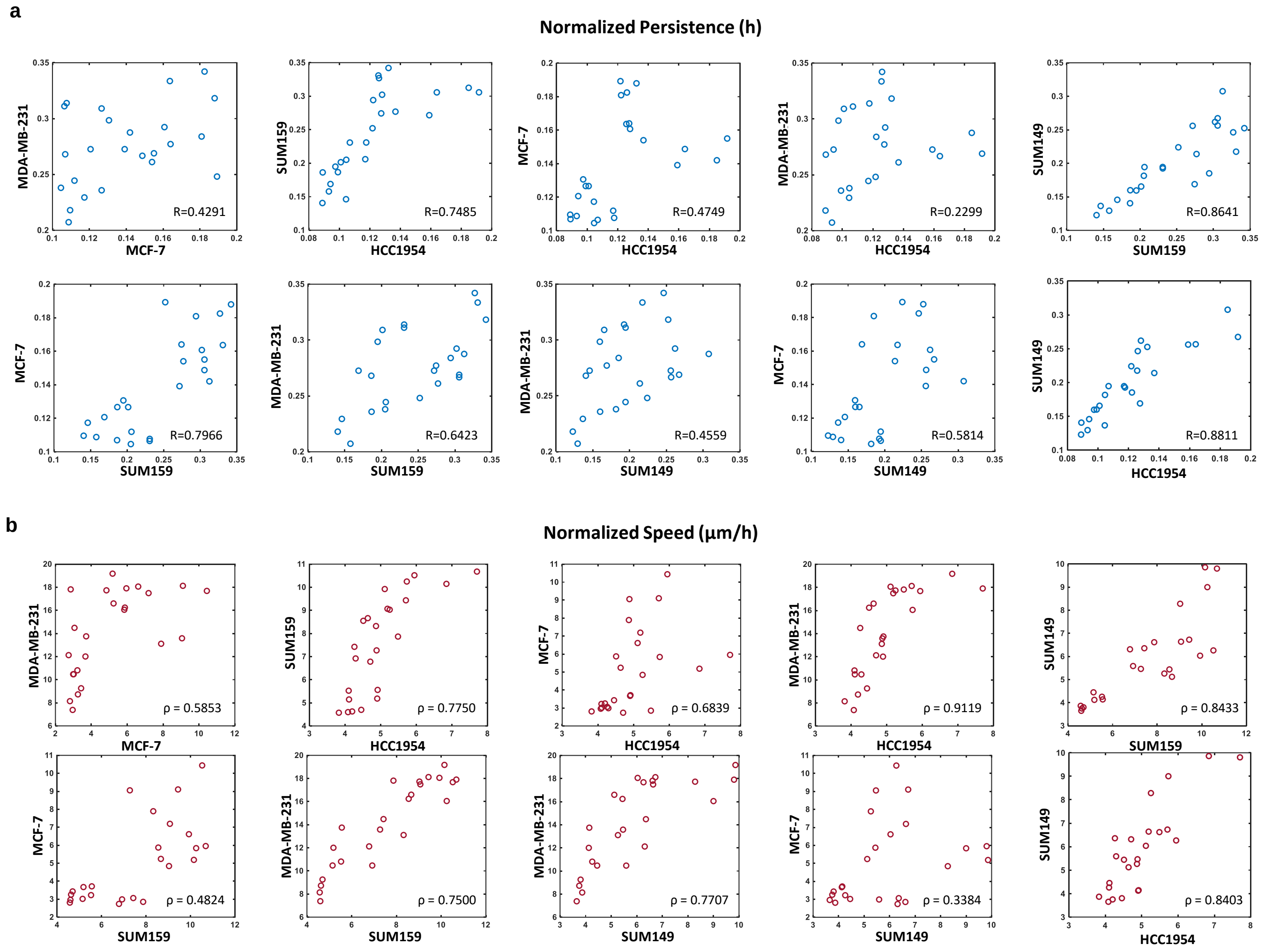


**Supplementary Figure 6. Correlation plots comparing the speed and persistence response to microenvironmental factors for all cell lines.** Comparison between all cell lines for normalized **(a)** speed and **(b)** persistence values in response to various combinations of microenvironmental stimuli. Correlation (ρ) values of the cell responses are noted on the plots.

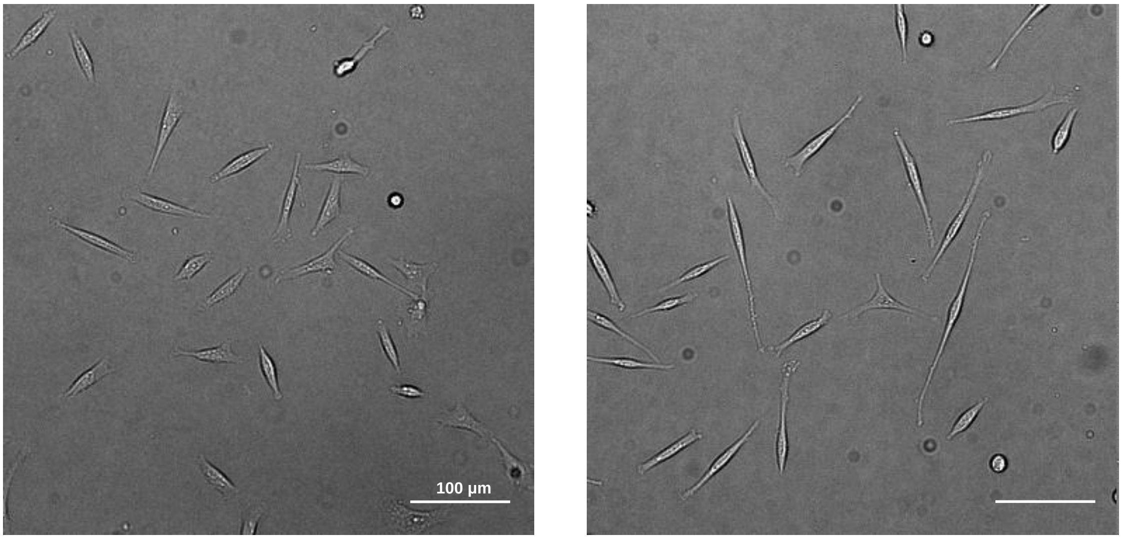


**Supplementary Figure 7. RHOA KO cells exhibited a morphological transformation compared to negative control MDA-MB-231 cells.** Representative brightfield images of **(a)** negative control and **(b)** RHOA KO MDA-MB-231 cells under the same microenvironmental conditions (0% FBS).

| | | | | | | | | | | | | | | | | |
|---|---|---|---|---|---|---|---|---|---|---|---|---|---|---|---|---|
| **Collagen** | - | - | - | - | - | - | - | - | + | + | + | + | + | + | + | + |
| **EGF** | - | - | + | + | - | - | + | + | - | - | + | + | - | - | + | + |
| **TNF-α** | - | - | - | - | + | + | + | + | - | - | - | - | + | + | + | + |
| **FBS (%)** | 0 | 10 | 0 | 10 | 0 | 10 | 0 | 10 | 0 | 10 | 0 | 10 | 0 | 10 | 0 | 10 |
| **RHOA KO** | 3.2 | 4.6 | 3.4 | 5.5 | 3.4 | 5.3 | 3.4 | 6.4 | 6.6 | 12.7 | 7.0 | 15.7 | 5.8 | 13.1 | 6.7 | 15.0 |
| **ARPC2 KO** | 3.3 | 4.1 | 3.5 | 4.4 | 3.1 | 3.9 | 3.3 | 4.5 | 5.5 | 8.5 | 6.8 | 11.0 | 5.2 | 8.1 | 6.1 | 9.8 |
| **CTTN KO** | 3.1 | 4.9 | 3.5 | 6.7 | 3.1 | 4.9 | 3.5 | 6.7 | 5.6 | 10.4 | 6.7 | 14.2 | 5.4 | 9.7 | 6.5 | 12.8 |
| **- CTRL** | 3.4 | 5.0 | 4.1 | 7.1 | 3.5 | 5.3 | 4.2 | 7.5 | 7.3 | 12.3 | 8.9 | 15.4 | 6.4 | 10.4 | 7.6 | 13.7 |

Speed (µm/h)

**Supplementary Figure 8. Heatmap summarizing the motility effects from the various combinations of microenvironmental factors on CRISPR KOs of MDA-MB-231.** Plot compares the motility of RHOA, ARPC2, and CTTN KOs to the negative control cell motility when under the effects of various combinations of microenvironmental factors. The microenvironmental factors included in this study are collagen (50 µg/mL), EGF (100 ng/mL), TNF- α (100 ng/mL) and FBS (0%, 10%).
